# Holsteins Favor Heifers, Not Bulls: Biased Milk Production Programmed during Pregnancy as a Function of Fetal Sex

**DOI:** 10.1101/002063

**Authors:** Hinde Katie, Carpenter Abigail J., Clay John S., Bradford Barry J.

## Abstract

Mammalian females pay high energetic costs for reproduction, the greatest of which is imposed by lactation. The synthesis of milk requires, in part, the mobilization of bodily reserves to nourish developing young. Numerous hypotheses have been advanced to predict how mothers will differentially invest in sons and daughters, however few studies have addressed sex-biased milk synthesis. Here we leverage the dairy cow model to investigate such phenomena. Using 2.39 million lactation records from 1.49 million dairy cows, we demonstrate that the sex of the fetus influences the capacity of the mammary gland to synthesize milk during lactation. Cows favor daughters, producing significantly more milk for daughters than for sons across lactation. Using a sub-sample of this dataset (N = 113,750 subjects) we further demonstrate that the effects of fetal sex interact dynamically across parities, whereby the sex of the fetus being gestated can enhance or diminish the production of milk during an established lactation. Moreover the sex of the fetus gestated on the first parity has persistent consequences for milk synthesis on the subsequent parity. Specifically, gestation of a daughter on the first parity increases milk production by ∼445 kg over the first two lactations. Our results identify a dramatic and sustained programming of mammary function by offspring *in utero*. Nutritional and endocrine conditions *in utero* are known to have pronounced and long-term effects on progeny, but the ways in which the progeny has sustained physiological effects on the dam have received little attention to date.

## INTRODUCTION

Since the 1970s, biologists have directed substantial research effort to understanding adaptive sex-biased allocation of maternal resources in animals and plants. Biologists have proposed numerous hypotheses for sex-biases, including local resource competition [1-2], “advantaged daughters” [3], local resource enhancement [4-5], the “safe bet”/reproductive value [6-7] and sex-differentiated sources of mortality [8]. The most well-known and investigated, though, remains the Trivers-Willard hypothesis [9]. Trivers and Willard hypothesized that a female, as a function of her condition, is expected to preferentially allocate resources to the sex that provides greater marginal return on that investment [9]. In polygynous mating systems characterized by male-male competition, they predicted that good condition females would bias resource allocation in favor of sons because males profit more form additional investment than do females [9]. Collectively, the hypotheses proposed in the literature can be loosely grouped according to the extent that the directionality of the sex-bias is contingent on maternal condition; however, the predictions deriving from these hypotheses are not always mutually exclusive, complicating interpretation of empirical results [10]. Large-bodied ungulates are frequently used for investigating sex-biased maternal allocation because male body size contributes substantially to success in competitive access to mating opportunities, but evidence for systematic sex-biases has been equivocal [10-14].

Although sex-ratio at birth has been the primary outcome investigated, post-natal maternal physiological transfer and behavioral care afford females substantial flexibility in sex-biased resource allocation [12]. Sex-biased nursing behavior has been investigated as a possible proxy for sex-biased milk production in numerous mammalian taxa [15-21]. Suckling behavior, however, is not useful for estimating milk energy transfer as verified by experimental use of radio-labeled isotopes in *Equus caballus* [22]. Direct evidence for sex-biased milk synthesis among non-domesticated species has now been reported in ungulates (*Cervus elaphus hispanicus*, [23]), rodents (*Myodes glareolus* [24]), primates (*Macaca mulatta* [25-26]; *Homo sapiens* [27-29], but see also [30] for exception), and marsupials (*Macropus eugenii,*[31]). Drawing systematic conclusions from the studies to date, however, is challenging in part because most have been limited by relatively small sample sizes or report milk composition without accounting for milk yield. The most comprehensive data derive from Iberian red deer (*Cervus elaphus hispanicus*) and rhesus macaques (*Macaca mulatta*). Landete-Castillejos and colleagues showed that hinds favored sons by producing more milk with higher protein content for them [23]. This bias did not vary as a function of maternal mass or age [23]. Among rhesus macaques, mothers produced higher milk energy density [kcal/g] for sons [26] due to higher fat content [25]. There was additionally an interaction with maternal life-history; smaller, younger mothers produced even higher fat and protein concentrations for sons and lower concentrations for daughters than did multiparous mothers [25]. However, at peak lactation, mothers of daughters, across parities, produced greater milk volume that offset the reduced energetic density of milk for daughters [26]. These studies failed to support sex-bias hypotheses that predict mothers in better condition will preferentially allocate resources to a particular sex, suggesting instead that there may be systematic sex-biases that are independent of maternal condition.

Mother’s milk, however, is particularly difficult to evaluate when investigating adaptive allocation of maternal resources. Milk synthesis is unlikely to be at the maternal optimum because of parent-offspring conflict [32-33]. Rather milk reflects a complex physiological and behavioral negotiation between the mother and offspring [34-35]. Functional development of the mammary gland initially occurs during pregnancy and is orchestrated by maternal and placental hormones, particularly placental lactogen, estrogen, and progesterone [36-38]. Post-natally, local regulation of milk synthesis is maintained by milk removal via offspring suckling [36,39] but maternal rejection can prevent or limit milk intake [18]. As a result, sex-biased milk synthesis may reflect differential cellular capacity in the mammary gland, programmed via hormonal signals from the fetal-placental unit, or post-natally through sex-biased nursing behavior [26]. There has been only one study that has investigated mechanisms underlying sex-biased milk synthesis. Koskela and colleagues used an elegant cross-fostering design in bank voles (*Myodes glareolus*) to demonstrate that all-female litters received significantly greater milk yield than did all-male litters, regardless of litter size or maternal condition [24]. The manipulation was conducted after females gave birth, and the extent to which pre-natal mammary gland development may have been sensitive to litter sex-ratio was not reported. Litter size during gestation has been shown to influence mammary gland development in sheep [40] and milk volume in goats [41], but the effect of fetal sex on milk synthesis has not been investigated.

We investigated the magnitude and direction of sex-biased milk synthesis in the Holstein breed of *Bos taurus.* Although intensive artificial selection has shaped cattle during recent centuries, domesticated cattle are derived from large-bodied, sexually-dimorphic aurochs (*Bos primigenius*) [42-43]. Among beef cattle, several small studies have revealed sex-biased milk production that favors sons [44], favors daughters [45], or no sex-biases [46]. In contrast, standardized husbandry practices, systematic milking procedures, detailed record-keeping, and large samples sizes make the dairy cow a powerful model for the exploration of maternal milk synthesis from both functional and mechanistic perspectives [35, 47–48]. Birth sex-ratio in dairy cows is male-biased [49], suggesting that mechanisms for sex-biases are operating in this taxon. Moreover the basic architecture for lactation is more highly conserved than other components of the genome, even for an animal artificially selected for milk yield [50]. Notably, because calves are removed from the dam within hours of parturition, this model system allowed us to investigate pre-natal mechanisms of sex-biased milk synthesis independent of post-natal maternal care and infant suckling behavior. Importantly, dairy cows are concurrently pregnant during lactation, typically 200+ days of the 305-day lactation [51]. We therefore predicted that milk synthesis on the first lactation could be affected not only by the sex of the calf produced, but also by the sex of the fetus gestated during lactation. We also predicted that mammary gland programming in response to fetal sex would persist into the subsequent lactation because the capacity to synthesize milk is, to some extent, cumulative across parities [52–54]. These complex predictions are clarified by schematic representation (Figure 1).

**Figure 1.**
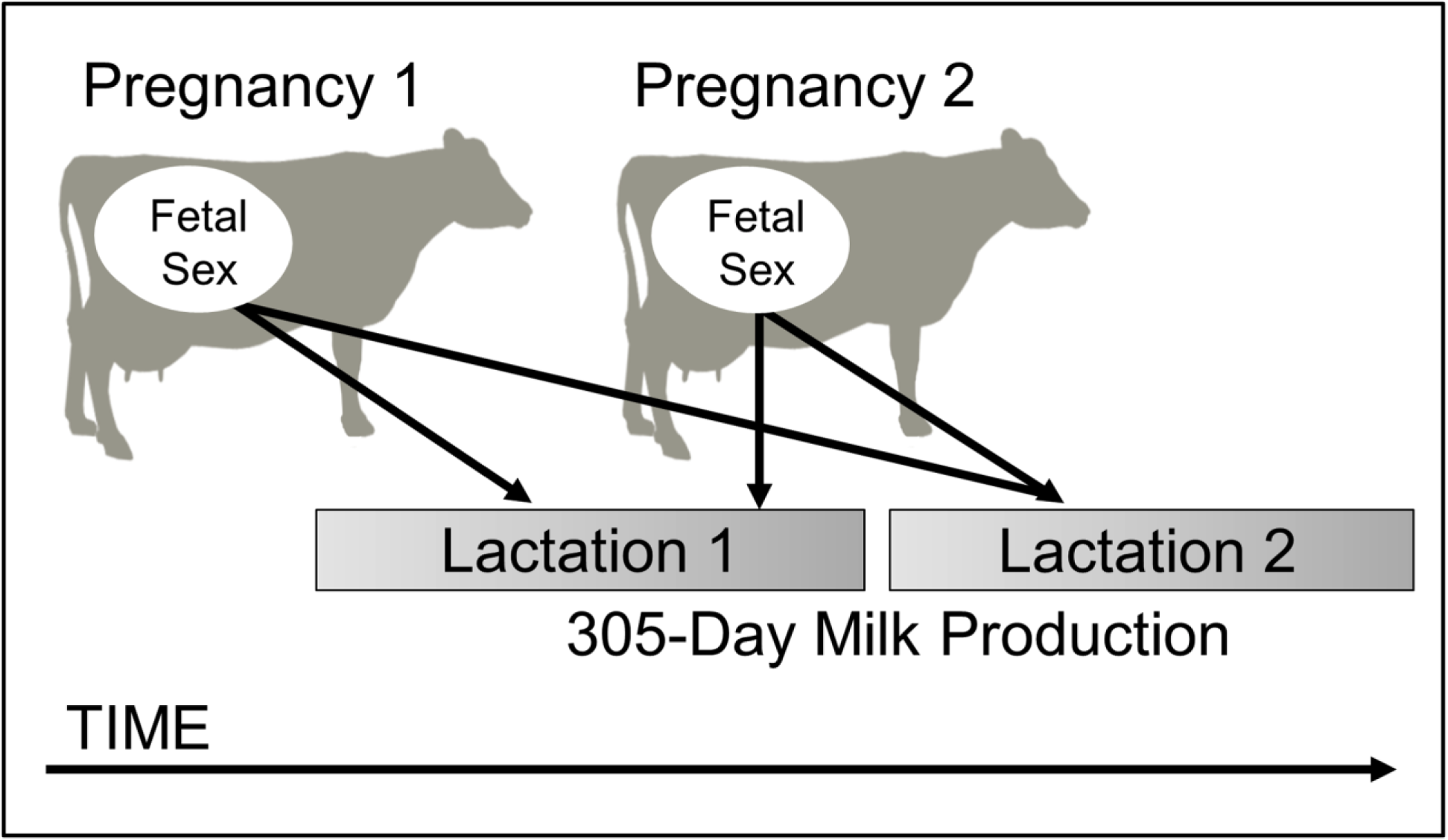
Hypothesis: milk production is influenced by fetal sex across lactations. Fetal sex in pregnancy 1 may alter milk production across multiple lactations because of the critical steps in mammary development that occur during the first pregnancy. In the cow, pregnancy 2 typically overlaps with lactation 1, providing opportunity for calf sex in parity 2 to impact milk production in the first lactation.

## METHODS

To investigate sex-biased milk synthesis, we acquired all lactation records from 1995 to 1999 in the database managed by Dairy Records Management Systems (http://www.drms.org). Whole-lactation milk yield and composition data were derived from monthly yield and composition data collected on commercial dairy farms across the United States. Standardized lactation curves, characterized over 5 decades of research, were then used to predict production between the monthly data points. Production is adjusted for breed, region, season and parity during the calculation of whole-lactation milk and component production, which was standardized to a 305-day lactation. These records are used daily by most of the 50,000 dairy farmers in the U.S. to make management decisions. Detailed discussions of the program and data analysis have been published elsewhere [55–56]. Data from the late 1990’s were used to avoid the influence of sex-selected semen in artificial breeding programs in the commercial dairy industry, which became common in the mid-2000’s [57–58]. Additionally, this period of time allowed for analysis of the effects of recombinant bovine somatotropin (bST) [59], approved in 1993 for commercial use in the U.S. The DRMS database includes a field for reporting administration of bST that was introduced into their software (PCDart) from the start of the commercial availability of bST.

Several steps were taken to clean the data prior to analysis. Only records from Holstein cattle were retained, and lactations that began with either twin births or abortions were excluded. Lactations with missing or corrupt lactation number, year, or calf sex designations were deleted. Duplicate records for a single lactation within cow were eliminated, and records for lactation ≥6 (representing 3.02% of lactations in the database) were excluded to enable repeated measures analysis of lactations with adequate representation in the database. If at least 1 of the first 5 test days, typically conducted monthly, were flagged for bST administration, then the lactation was considered bST-positive (N = 100,478; 3.9% of lactations). The final database consisted of 2.39 million lactation records, representing 1.49 million individual Holstein cows, however due to missing data in certain fields, some analyses included fewer lactations and final analysis sample sizes are reported for each analysis. Mixed models were used to evaluate the fixed effects of calf sex, parity, bST, and interactions and the random effect of year according to the following model:

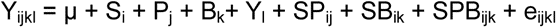

where Y_ijkl_ is a dependent variable, µ is the overall mean, S_i_ is the fixed effect of calf sex (i = 1 to 2), P_j_ is the fixed effect of parity (j = 1 to 5), B_k_ is the fixed effect of bST (k = 1 to 2), Y_l_ is the random effect of year (l = 1 to 5), SP_ij_ is the interaction of calf sex and parity, SB_ik_ is the interaction of calf sex and bST, SPB_ijk_ is the interaction of calf sex, parity, and bST, and e_ijkl_ is the residual error. Repeated lactations within cow were fit to a heterogeneous autoregressive (ARH[1]) covariance structure. Analyses were completed using the Mixed Procedure of SAS (version 9.3; SAS Institute, Cary, NC). Significant interactions were investigated using the SLICE option and means were separated using the PDIFF option of SAS, with significance declared at *P* < 0.05.

To exclude potentially confounding effects of dystocia and bST treatment on results and to evaluate carryover effects of calf sex on multiple lactations, a more conservative data set was generated. All bST-positive lactations were deleted, and only those beginning with a calving difficulty score of 1 or 2 (no or minimal difficulty) were retained. Finally, the data were narrowed to only those cows with both lactations 1 and 2 represented, leaving 113,750 cows. Data for 305-day milk yield in lactations 1 and 2 were modeled with the fixed effects of calf sex_1_, calf sex_2_, calf sex_1_ × calf sex_2_, and year. Analyses were completed using the Mixed Procedure of SAS (SAS Institute) and means were separated using the PDIFF option of SAS, with significance declared at *P* < 0.05.

## RESULTS

### Sex-Biased Milk Synthesis: Full Dataset

Holsteins biased milk production in favor of daughters, producing significantly more milk over the 305 days of standard lactation after gestating a daughter (Fig. 2). These findings are based on 2.39 million lactation records from approximately 1.49 million female cows. First-parity cows giving birth to a daughter produced 142 ± 5.4 kg more milk over the 305-day lactation period than did those giving birth to a son (7,612 vs. 7,470 ± 69 kg, *P* < 0.001). Similar, though marginally smaller, effects were observed in parities 2–5 (Fig. 2A). The overall effect amounted to a 1.3% increase in whole-lactation milk production for cows bearing daughters (Table I). Extrapolation from total lactation production values revealed that milk composition was similar after gestation of a son or daughter. Fat concentration was 3.61% after gestation of a daughter and 3.62% after gestation of a son; protein concentrations were the same (3.17%).

**Figure 2.**
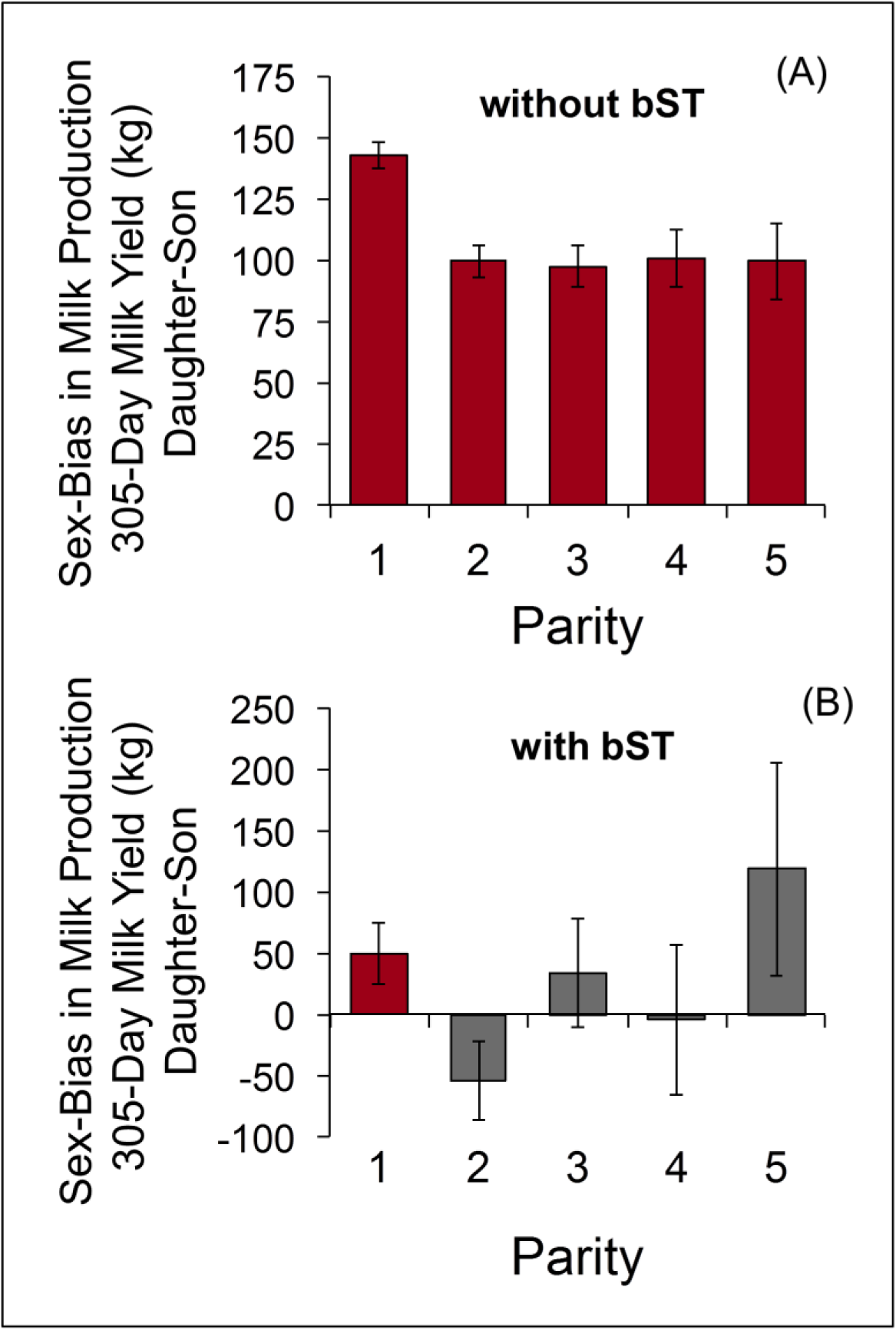
Daughters result in greater lactation productivity, and this effect is altered by exogenous somatotropin (bST) administration. Lactation records from Holstein cows (N = 2.39 million lactations) were analyzed to determine effects of calf sex, parity, use of bST, and their interactions on 305-day milk production. Calf sex influence on milk production was dependent on bST use (interaction *P* < 0.01). A) In the absence of bST, daughters resulted in significantly greater milk production compared to sons across all parities (all *P* < 0.001). B) Lactations influenced by bST administration failed to consistently demonstrate the daughter bias. Daughters still conferred an advantage in first-parity cows administered bST (*P* < 0.05), but did not significantly influence milk yield in parity 2–5 cows. Values are differences of LS means ± SED.

**Table I.** Influence of calf sex, in the presence and absence of exogenous somatotropin (bST), on lactation productivity.

|  | No bST |  |  | bST |  |  |  | P value |  |  |
| --- | --- | --- | --- | --- | --- | --- | --- | --- | --- | --- |
| **Cow Milk Production** | **Daughter** | **Son** | **SEM** | **Daughter** | **Son** | **SEM** | **N (lactations)** | **Sex** | **bST** | **Interaction** |
| 305-day milk yield (kg) | 8172.8 | 8065 | 68.6 | 9123.4 | 9094.8 | 70.8 | 2,391,300 | < 0.001 | < 0.001 | < 0.01 |
| 305-day fat yield (kg) | 295.56 | 291.5 | 3.25 | 329.5 | 328.11 | 3.4 | 2,125,643 | < 0.001 | < 0.001 | < 0.01 |
| 305-day protein yield (kg) | 258.78 | 255.6 | 2.06 | 291.05 | 290.26 | 2.16 | 2,108,796 | < 0.001 | < 0.001 | < 0.01 |
| Peak milk (kg/d) | 36.97 | 36.36 | 0.34 | 40.52 | 40.38 | 0.35 | 825,175 | < 0.001 | < 0.001 | < 0.01 |

The disparity between milk produced following birth of a son vs. a daughter was largely eliminated by the use of bST. A recombinant, exogenous form of the growth hormone somatotropin, bST promotes endocrine alterations to partition a greater proportion of nutrient supply to the mammary gland, thereby increasing milk production [60]. Recombinant bST is approved for exogenous administration to dairy cows beginning at week 9 of lactation. In our sample, bST accounted for a 12% increase in whole-lactation milk yield (Table I). On first parity, cows administered bST still produced significantly higher milk yield if they had a daughter (8,681 vs. 8,631 ± 71 kg, *P* < 0.05), but sex-biased milk synthesis was not observed in parities 2-5 (Fig. 2B).

### Sex-Biased Milk Synthesis: Conservative Sample

Male calves are typically larger than females, and pose a greater risk of dystocia [61–62]. Dystocia is associated with decreases in whole-lactation milk production [62], and we hypothesized that the milk yield advantage conferred by a daughter might have been at least partly due to decreased incidence of dystocia compared to delivery of sons. Indeed, in our data, the odds of a son inducing dystocia (calving difficulty score ≥ 3 on a scale of 1 to 5) were significantly greater than for daughters (5.6 vs. 4.2% incidence, *P* < 0.001, odds ratio 95% CI: 1.32–1.35). Nevertheless, sex-biased milk synthesis remained when analysis was restricted to a subset of the dataset (N = 113,750) that excluded cases of bST and dystocia, and included information on individual cows across the first and second parity. On first parity, cows producing daughters had significantly greater 305-day milk yield, with an advantage of 1.6% relative to cows producing sons (7,947 vs. 7,818 ± 9.6 kg, *P* < 0.001). The daughter advantage was also observed in parity 2, although the magnitude of the difference was reduced (0.83%; 8,515 vs. 8,445 ± 37 kg, *P* < 0.001). These results indicate that the milk production advantage associated with birth of a daughter is not attributable to prevention of dystocia.

### Inter-Parity Consequences of Fetal Sex

Milk production on first lactation was associated with the sex of the fetus on the second pregnancy because the two overlapped temporally (Figure 3A). Across the first two parities in the subset that excluded cases of bST and dystocia, birth combinations could be son_1_son_2_, son_1_daughter_2_, daughter_1_son_2_, and daughter_1_daughter_2_. Cows that had first produced a son and were gestating a son for their second pregnancy synthesized significantly less milk over 305 days than did all other groups (*P* < 0.001; son_1_son_2_ = 7,768 ± 11.4 kg, N = 32,294). Gestation of a daughter on the second pregnancy could partially “rescue” milk synthesis on the first lactation if a son had been produced previously (*P* < 0.001; son_1_daughter_2_ = 7,876 ± 12.2 kg, N = 27,807), but remained significantly less than cows that had produced a daughter on their first pregnancy (*P* < 0.001). Fetal sex on the second pregnancy didn’t have any effect for cows that produced a daughter on pregnancy 1 (daughter_1_son_2_ and daughter_1_daughter_2_ were 7,940 ± 12.3 kg, N = 27,834 and 7,954 ± 12.6 kg, N = 25,815, respectively; *P* = 0.36).

**Figure 3.**
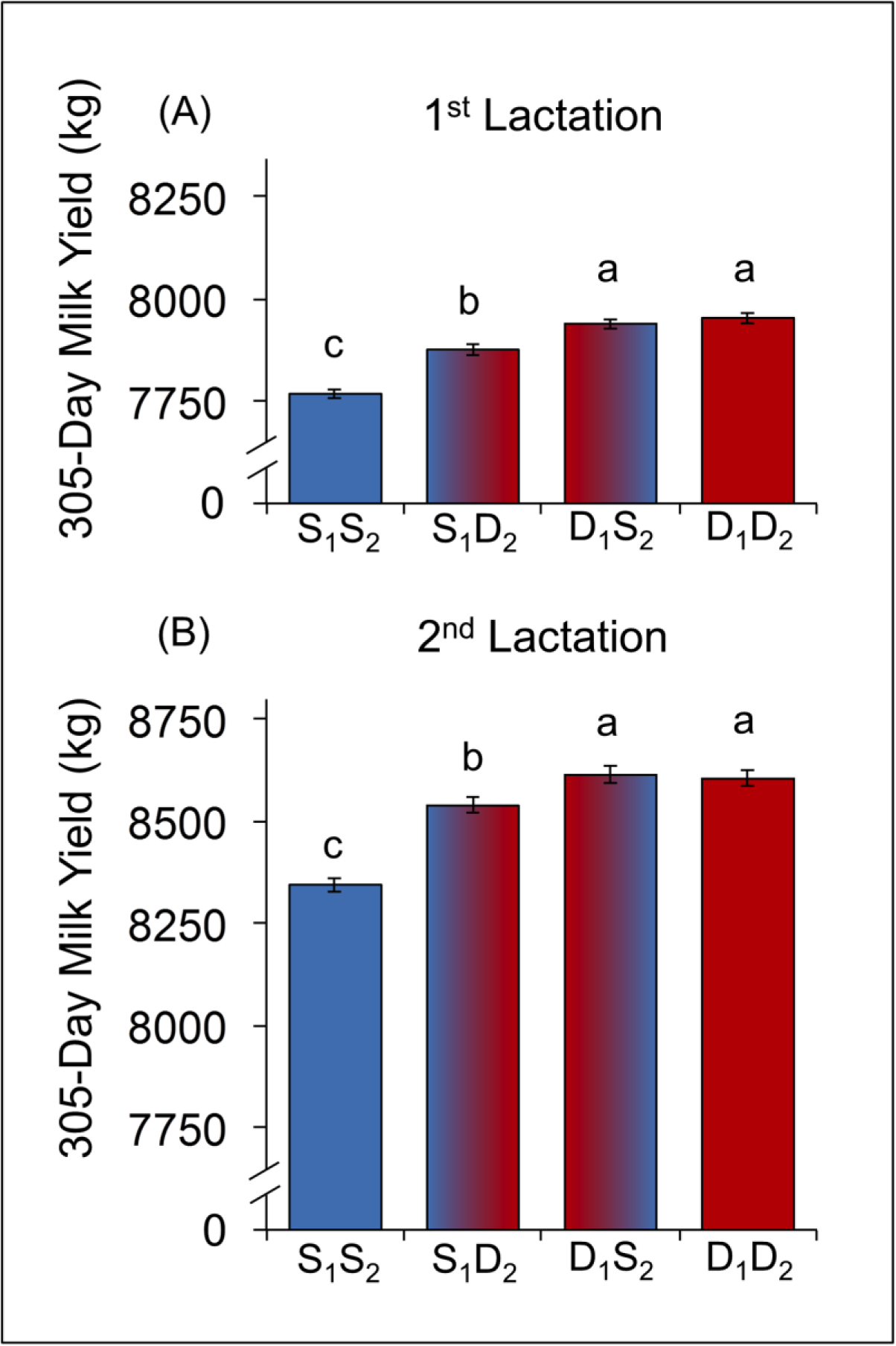
Daughters confer milk production advantages post-natally, during gestation, and across multiple lactations. Cows (n = 113,750) with both first and second parity lactation records, with no reports of dystocia or bST administration, were used to assess effects of calf sex on milk production in the first 2 lactations. Groups are labeled by calf sex (S = son, D = daughter), with the pregnancy denoted by subscript. Values are LS means ± SEM. Means labeled with different letters differ (*P* < 0.001), and those with common labels do not (*P* > 0.10). A) First-parity cows having a daughter produced significantly more milk than those having a son, but gestating a daughter in pregnancy 2 increased milk production in cows that had a son first. B) Second-parity milk production is greatest in cows that had a daughter in pregnancy 1. Additionally, cows with a son in pregnancy 1 showed increased milk production if they had a daughter in pregnancy 2.

Fetal sex on the first parity had persistent effects on milk production during the second lactation (Figure 3B). Cows that produced a son on their first parity were handicapped in their milk production on their second lactation (*P* < 0.001), particularly if they gestated a son on the second pregnancy as well (son_1_son_2_ = 8,345 ± 18.9 kg). Production of a daughter on the second parity partially increased milk production on second lactation (*P* < 0.001; son_1_daughter_2_ = 8,539 ± 19.4 kg). Cows that produced a daughter on their first parity produced significantly more milk on their second lactation (*P* < 0.001), regardless of the sex of the calf on the second parity (daughter_1_son_2_ and daughter_1_daughter_2_ were 8,614 ± 19.6 kg and 8,605 ± 19.8 kg, respectively; *P* = 0.19).

## DISCUSSION

Holstein dairy cows demonstrate a significant biological effect of sex-biased milk production in favor of daughters. In dairying, calves are removed on the day of birth and standardized mechanical procedures are used for milking, therefore post-natal sex-bias does not explain the results presented here. Instead milk production varied as a function of fetal sex, indicating that functional development of the mammary gland is influenced pre-natally. Importantly, lower milk yield for sons was not compensated by higher protein and fat production; total production of milk energy was greater in cows that gestated daughters. Among rhesus monkeys, mothers rearing daughters produce more milk, but of significantly lower milk energy density-the aggregated calories derived from fat, protein, and sugar-than do mothers of sons [26]. To our knowledge, the results reported here are the first to document that fetal sex influences milk production. Moreover the effects on milk production were dynamic and persistent across parities. Importantly, gestation of a daughter on the first parity increased milk production across the first two lactations and was protective against the negative effects of male gestation on the second parity. In contrast, gestating a son on the first parity suppressed milk production on the first two lactations, but the conception of a daughter on the second parity partially improved milk production. Nutritional and endocrine conditions *in utero* are known to have pronounced and long-term effects on progeny [63], but the ways in which the progeny has sustained physiological effects on the dam have been less studied.

Sex-differentiated programming of the mammary gland is further substantiated by the greater effect of bST administration in cows gestating sons than cows gestating daughters. Postnatal administration of recombinant bovine somatotropin (bST) in multiparous cows overwhelmed the prenatal effects of offspring sex, but had a greater effect in cows gestating sons. Somatotropin, or growth hormone (GH), is produced in the anterior pituitary, stimulated by GH-releasing hormone. Most notably, GH influences metabolism in hepatic and adipose tissues, shunting more maternal bodily reserves to milk synthesis [64]. Insulin-like growth factors are believed to be the major mediators of the effect of GH on the mammary gland [60], however GH also directly affects the mammary gland and increases milk synthesis [65–66]. While the mean production parameters increased with the administration of bST for cows producing both daughters and sons, the proportional increase in milk production was greater for multiparous cows gestating sons. Rose and colleagues reported that cows that had low milk yield responses to bST treatment within a herd had greater milk yields before bST treatment compared to cows with a high response in milk yield [67]. This is consistent with our results that cows birthing daughters had elevated milk production and a lower response to exogenous bST administration compared to their counterparts bearing sons. We posit that mechanisms underlying lower initial milk production and greater individual response to bST administration are likely responsible for the greater response to bST in cows with sons. Administration of bST in many ways represents an “experimental” manipulation of mammary gland programming and reveals possible mechanistic pathways through which sex-biases are operating. Although bST was able to overwhelm sex-biased milk synthesis among multiparous cows, significant sex-bias remained among primiparous cows whose mammary glands had functionally developed for the first time in the context of the fetal sex of the first gestation. The magnitude of sex bias is strongest among first parity rhesus monkeys [25–26] and possibly humans [28–29] and Tamar wallabies [31] in which primiparous females have been disproportionately represented in published studies. The effect of fetal sex may diminish to some extent among multiparous females due to the aggregate effects on mammary gland architecture of sequential gestations of different fetal sexes. Alternatively, maternal investment tactics may change as a function of residual reproductive value [68] or targeted effort during critical developmental windows [69].

These biological findings may have economic impact for the modern dairy industry. With the widespread availability of sexed-selected semen for use in artificial breeding programs, dairy managers have the option of achieving approximately 90% female pregnancies rather than a natural rate near 47% [49]. There are many factors for managers to consider when evaluating the profitability of sexed semen use, including decreased conception rate [57] and increased semen cost. Some published analyses have been skeptical of the economic merit of using sexed semen on dairy operations [70], although the cost of the cell sorting technology continues to drop, making recent analyses more favorable [71]. Accounting for the impact of a female calf on lactation productivity revealed by our analysis, however, further improves the expected profitability of sexed semen use. It is common to use sexed semen for breeding nulliparous heifers only, and given the long-term impact of a first-parity daughter, the production benefits of this management strategy are substantial. The cumulative increase in milk yield over two lactations for a cow giving birth to a daughter on the first parity rather than consecutive bulls is ∼445 kg (Fig. 3). The impact of sexed semen on the structure of the dairy industry has been a complex question already [72], but these results highlight a key factor that has not previously been considered.

The precise mechanistic pathways through which fetal sex influences mammary gland development remain unknown. Fetal-origin hormones may translocate via maternal circulation to bind directly to receptors in the dam’s mammary gland influencing functional development and subsequent milk synthesis. Among ungulates, ruminants may be especially valuable for understanding mammary gland development during pregnancy as a function of fetal sex because of their cotyledonary placenta. Klisch and Mess posited that for ruminants, an evolutionary “arms race” between the mother and fetus [73] for glucose transport, necessitated by the lack of gastrointestinal glucose supply [74], resulted in selective pressure that favored an “inefficient” placenta [75]. For example, the placenta of the domestic cow has ∼5 times the surface area as the horse placenta even though the two species produce similarly sized neonates [76]. As a byproduct of the greater placental surface area, fetal steroidal hormones can readily diffuse into maternal circulation [75]. Concentrations of estrogens and androgens differ between male and female fetuses and, if in maternal circulation, potentially enhance or inhibit mammary gland development and consequently milk synthesis during lactation. In dairy cows, fetal steroid hormones are present from the first trimester and are critical for the development of fetal sex organs [77–78]. Insulin-like peptide 3 (INSL3), another fetal-origin bioactive, increases in maternal circulation across pregnancy in dairy cows gestating sons and decreases in cows gestating daughters [79] but the influences of fetal-origin INSL3 on the mammary gland are not known. Functional development of the mammary gland in taxa characterized by highly invasive hemochorial placentas may also be susceptible to fetal hormones; indeed the majority of reports of sex-biased milk synthesis in the literature are from taxa that have greater placental invasion and/or placental surface area [63, 76, 80]. Suggestively, human mothers with higher concentrations of circulating androgens during the 2^nd^ trimester had a lower probability of sustaining breastfeeding to three months post-partum [81]. The higher circulating androgens may have originated from fetal sons, but the effect of fetal sex was not directly analyzed in that study, nor was milk synthesis measured. Indirectly, fetal sex may influence the production of placental lactogen, a primary hormonal driver of mammary gland development during pregnancy [36–38] but as of yet differences in placental lactogen as a function of fetal sex have not been reported.

Daughter-biased milk synthesis may reflect adaptive maternal allocation in response to fetal sex, adaptive fetal manipulation of maternal physiology, or may be a by-product of the maternal-fetal interface. Importantly, uniformly biased milk production in favor of daughters across maternal conditions does not support the Trivers-Willard hypothesis [9], or other hypotheses positing facultative sex-biased allocation of resources as a function of maternal condition [10]. Dairy cows have a male-biased birth ratio; in the absence of sex-specific artificial insemination, between 50-54% of calves born are male [49, 82]. The mediating effect of maternal condition on birth-sex ratio has been inconsistent [83] as has been the directionality of birth sex-ratio bias. Better-condition cows may produce more sons [84] or daughters [85]. Integrating the results presented here, dairy cows produce more sons, but seemingly favor daughters with more milk. Mammalian mothers in polygynous taxa may preferentially allocate physiological resources to daughters so that they are able to initiate reproduction at relatively younger ages than do sons [26, 86]. For female mammals, because of the temporal constraints of pregnancy and lactation, lifetime reproductive success of daughters will be contingent on the length of their reproductive careers, a function of age at first birth and longevity [87–88]. Among sexually dimorphic polygynous taxa, the temporal constraints are relaxed for males, who benefit from growing bigger and stronger [89–90], allowing males more time to compensate for deficits in early life maternal investment before becoming reproductively active [91]. Daughter-biased milk production may involve life-history tradeoffs for both cows and their daughters. High milk production in dairy cows is generally associated with reduced fertility, health, and survival depending on environmental conditions [92]. Moreover daughters gestated during lactation have moderately reduced survival and milk production in their own adulthood [93–94]. Although we do not know whether the magnitude of the effects presented here is correlated with such consequences, future research should investigate the fitness effects of daughter-biased milk synthesis both in the short-term (i.e. inter-birth interval), across the lifetime, and inter-generationally.

The question remains though, under natural conditions how do bull calves grow faster during the post-natal period if their dams are producing less milk, and therefore lower total protein and fat production? One explanation may be that females bias nursing behavior such that milk production is up-regulated for sons, a tactic we could not evaluate in conventional dairying as calves are removed after birth. Landete-Castillejos and colleagues revealed that among captive Iberian red deer, dams rearing sons had greater total milk production and total protein production [23], possibly due to post-natal hind-calf behavioral dynamics. However in the one study to date of cow maternal behavior, cows do not show any sex biases in nursing behavior [18]. In beef cattle that are reared by their dam, sons are born bigger and have better post-natal growth than do daughters, but only one out of three studies has shown any evidence of male-biased milk synthesis [44–46]. In the absence of post-natal behavioral modifications of prenatal mammary gland programming, the presence and concentration of other milk bioactives such as immunofactors and hormones that influence offspring development [35] may differ in milk produced for sons and daughters. Notably, investigations of sexually dimorphic developmental trajectories, however, overwhelmingly essentialize the role of the mother and sex-biased allocation of maternal resources. More often overlooked are sexually differentiated mechanisms within offspring that influence utilization and assimilation of early life nutrition and environmental signals [26, 95–96]. Consideration of progeny-specific adaptations as well as biased maternal effort will contribute to a better understanding of the ontogeny of sexual dimorphism.

## Acknowledgements

We thank Joan Silk, Julienne Rutherford, and two anonymous reviewers for their comments on earlier drafts of the manuscript. Contribution no. 13-391-J from the Kansas Agricultural Experiment Station (BJB and AJC). The funders had no role in study design, data collection and analysis, decision to publish, or preparation of the manuscript. No additional external funding was received for this study.

